# Exogenous pigments shield microorganisms from spaceflight-induced changes

**DOI:** 10.1101/2021.07.29.454367

**Authors:** S. Sharma, R. S. H. Smith, N. A. Lee, S. L. Wilson, M. M. Smith, N. Oxman

## Abstract

Research has indicated that pigments commonly produced by microorganisms may be protective against the environmental stresses inherent to spaceflight. However, few studies have directly tested the protective capabilities of microbial pigments applied externally as shielding materials. In this study, liquid cultures of *Bacillus subtilis* were shielded by various pigment solutions, and solid media cultures of *Bacillus subtilis* were co-inoculated with the highly pigmented microorganisms *Aspergillus niger* and *Neurospora crassa*. These experiments were conducted in a compact, automated payload aboard the International Space Station (ISS) interior for 30 days. Post-flight phenotypic analyses of liquid cultures showed that solutions of carotenoid pigments were effective at minimizing detrimental effects of spaceflight. Elevated growth rate was observed for solid cultures, and distinct morphology changes were identified in both liquid and solid samples and quantified as markers of spaceflight-induced stress. These findings collectively progress our understanding of microbial pigments for the development of space-related applications.

## Introduction

A primary aim of microbiological research in space is to assess the environmental factors relevant to life in upcoming manned missions^1^. Environmental cataloguing and experimental findings have demonstrated that spacecraft-associated microbiota and studied model organisms survive and adapt in the presence of multiple environmental stressors associated with space habitats, motivating investigation regarding the mechanisms behind their resilience in harsh conditions^7^. Particular emphasis has been placed on how microorganisms overcome or mitigate ionizing radiation—one of the greatest concerns for space missions^8^ —which has detrimental effects not only on living organisms, but also on electronic equipment and other materials^9^. The study of melanized organisms identified within microbial surveys of the International Space Station (ISS) has opened the possibility that melanin pigments may be protective in the space environment^10,11^.

Pigmentation is a potential strategy employed by microorganisms to survive in the presence of elevated ionizing radiation. ^12,13^. This capability, in addition to the known role of melanin in protecting from UV radiation^14,15^, has been investigated in highly pigmented fungi in a damaged Chernobyl nuclear reactor in 2000^12,13^ and has since led to the proposal of microbial melanins as a natural radioprotective material relevant for space travel^11,16^. A small number of studies have recently been conducted or are currently underway to evaluate melanin’s feasibility within living systems^17^ or as a material embedded within a polymer^18^. Other pigments, including carotenoids^19–21^, have also been demonstrated to have radioprotective capabilities and are found in extremophilic microorganisms such as the astrobiological model *Deinococcus radiodurans*^*22–24*^. They have been of interest in microbial biotechnology, yet as of now, have not been proposed as space-relevant protective materials nor thoroughly studied for similar applications. The study of pigments for both UV and ionizing radiation protection is a nascent field that warrants significant investigation given the known properties of melanin and carotenoids and their potential to be created on-site.

In this study, we investigated pigments as a class of biocompatible shield materials for protection from stressors in the spaceflight environment and examined their utility in a 30-day experiment aboard the ISS. The stressors within built habitats in a Low Earth Orbit (LEO) environment include ionizing radiation, microgravity^25^, and limited oxygen availability^26 27^. They have been demonstrated to affect microbial stress response, membrane-associated readjustments, metabolic rearrangements, and genetic alteration, which in turn affect survival^28^. We included multiple types of melanins and carotenoids as potential shields, and studied them both within natural systems (in vivo) and as externally applied liquid solutions (ex vivo). We evaluated the efficacy of these shields through the observation and quantification of phenotypic changes between spaceflight and ground control cultures. Our results indicate that exogenous pigments mitigate the impact of spaceflight environments as measured through phenotypic changes, including (i) growth dynamics and (ii) viable cell count in *B. subtilis*. We also identified two morphology metrics that are impacted by spaceflight: (i) novel morphotypes of *B. subtilis* that persist through multiple generations and (ii) irregular colony morphology in two different microorganisms exposed to spaceflight. In light of the increasing accessibility of ISS small experiment slots and development of automated payload platforms, the visible phenotypes and markers catalogued here form a promising approach by which future studies may continuously measure microbial adaptations with embedded camera systems inside miniaturized research modules during flight. Our findings indicate that pigments have potential as biocompatible shield materials in spaceflight. The data and approaches presented here provide a foundation for future applications-focused studies.

## Mission overview

The *Radiofungi* experiment was an autonomous payload within the NanoRacks BlackBox system that was housed within the interior environment of the International Space Station (ISS) in March 2020 for a period of 30 days (Fig. 1a). The payload enabled the examination of biophysical relationships between cells and their local environments, which are altered by microgravity and spaceflight^29^. Five microorganisms were chosen: *Aspergillus nige*r, a model fungus for melanin pigment production and spaceflight adaptation^30–32^; *Neurospora crassa*, a model filamentous fungus that produces carotenoids and has rarely been studied in spaceflight^33–35^; *Xanthophyllomyces dendrorhous*, a carotenoid-producing fungus^36–38^; *Rhizobium etli*, a model soil bacterium that produces melanin^39,40^; and *Bacillus subtilis*, a model bacterium commonly used in spaceflight research^29,41–44^. The following pigments were selected to provide a panel of potential shield materials: beta-carotene, lycopene, lutein, astaxanthin, eumelanin from *Sepia officinalis*, and synthetic eumelanin. Solutions of non-pigmented chemicals, L-tyrosine (a precursor to melanin) and amino acid L-cysteine, were included as controls to assess the effect of other aromatic and non-aromatic organic molecules.

**Figure 1.**
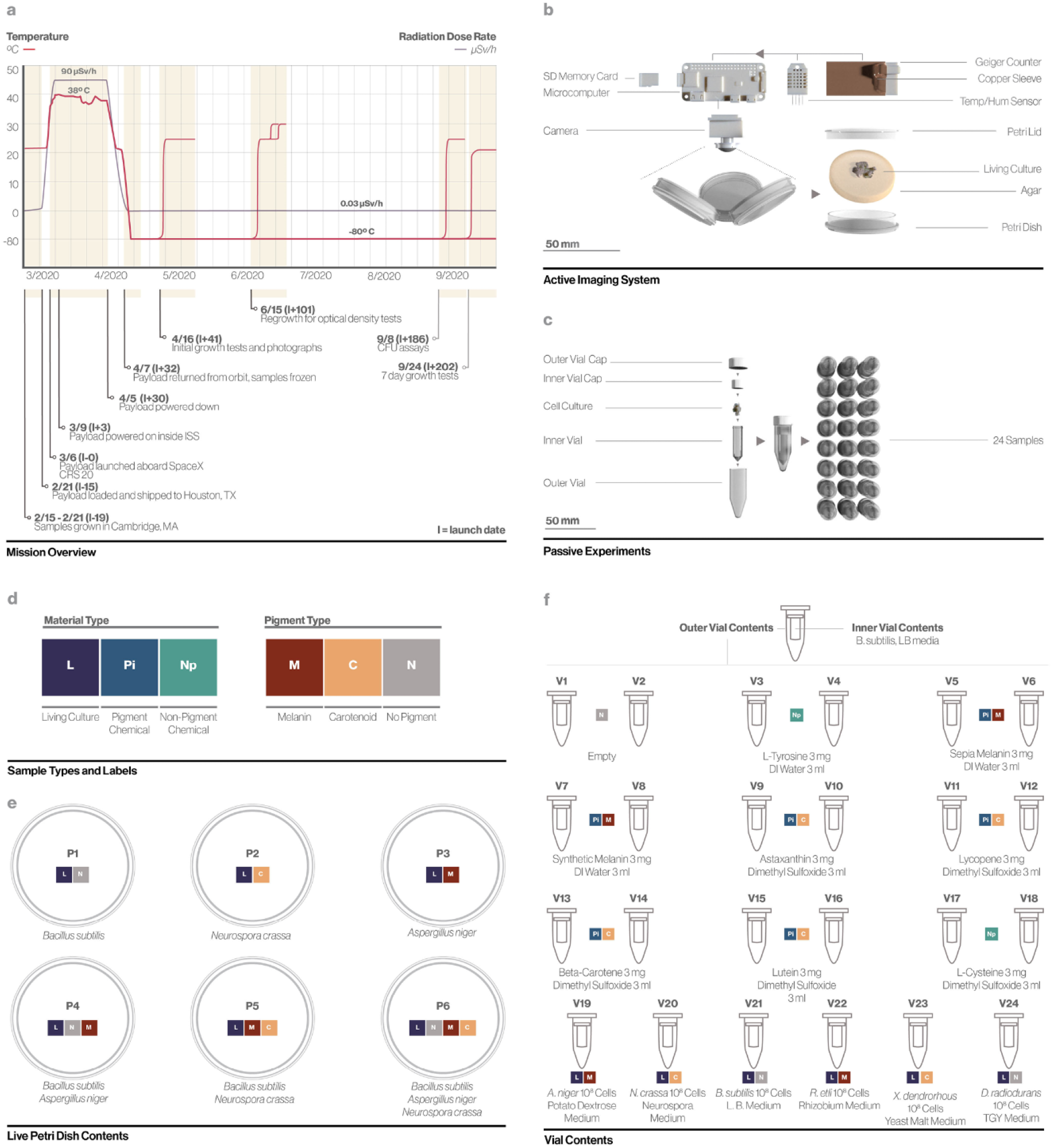
Overview of Radiofungi payload. (a) Mission overview with key dates, recorded temperature, and radiation dose rate. (b) Active payload components for recording temperature, humidity, radiation dose rate, and time-lapse photography. (c) Passive experiment components with two levels of containment. (d) Color labels for material and pigment type. (e) Labeled Petri dish sample contents. (f) Labeled vials for passive experiments with liquid B.subtilis liquid culture in inner vials and various shield solutions in outer vials.

The optimized configuration of the payload allowed for containment and inclusion of biological samples as well as an active environmental monitoring and imaging system (Supp. Fig. 1). Active and passive elements of the payload are presented in Fig. 1b and 1c, respectively. The active elements included six cultures of living microorganisms on hydrogel medium, hereafter described as “solid cultures”. These were imaged using two cameras at 10-minute intervals throughout the duration of the experimental period. The passive elements were 24 cultures of *B. subtilis* in 2mL vials, hereafter termed “liquid cultures”, placed within 5mL vials containing various solutions, hereafter described as “shields”. The components of the solid cultures are shown in Fig. 1e; the components of liquid cultures are shown in Fig. 1f.

## Results

### Autonomous environmental monitoring

The payload sensors for temperature, humidity, and radiation monitoring recorded continuously for the duration of spaceflight. The payload experienced 26 distinct and regular cycles of radiation at a rate similar to the expected dose rate for an unshielded object^45^ (Fig. 2a). The data were then analyzed with respect to the predicted orbital position of the ISS, revealing a correlation between dose rate and position (Fig. 2c). The payload experienced elevated temperature and humidity above those observed during pre-flight testing and validation, but these parameters remained within a realistic range for the dynamic target environment (Fig. 2b). This heat likely contributed to the hydrogel shrinkage and resulting humidity.

**Figure 2.**
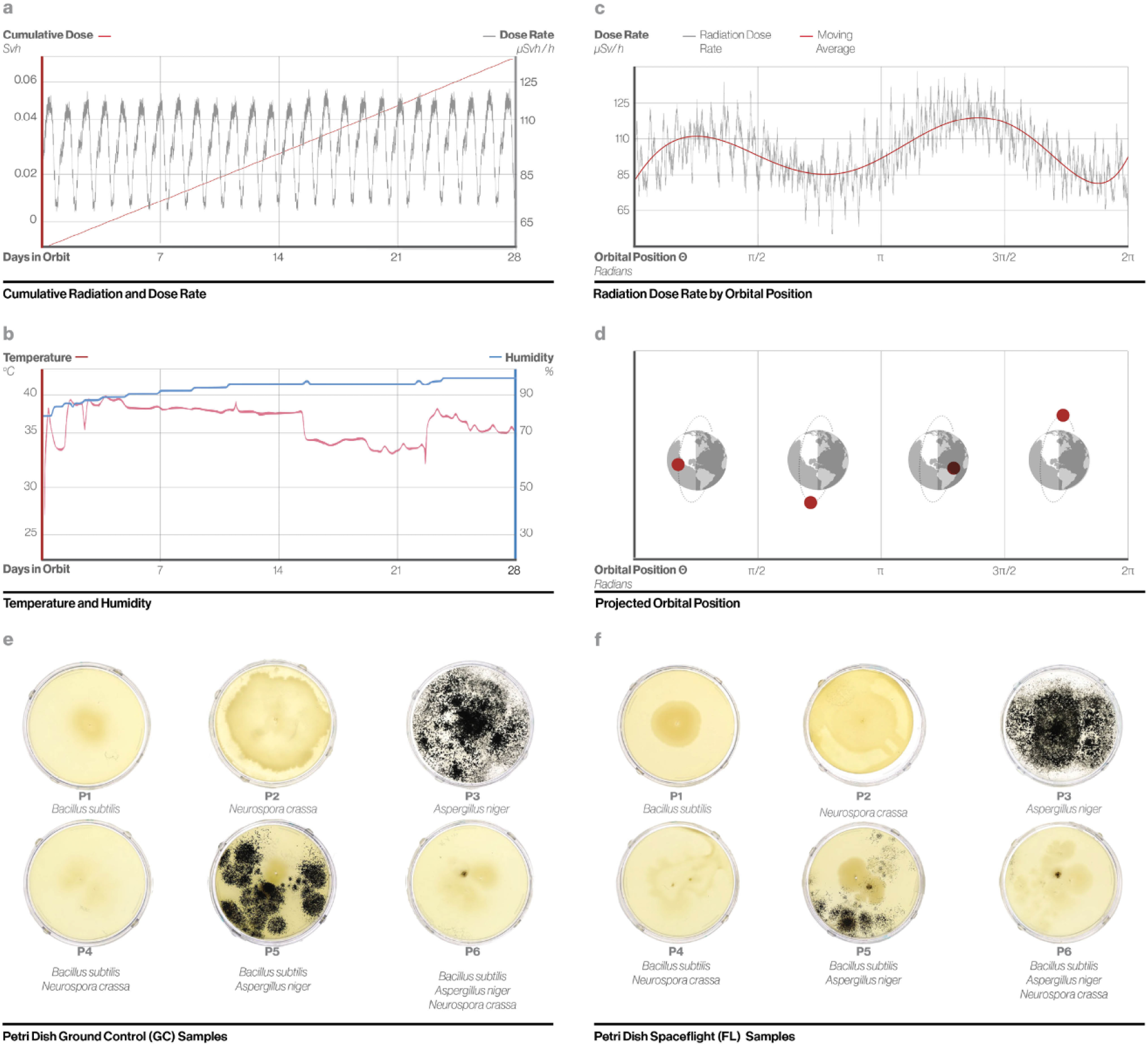
Autonomous payload environmental monitoring. (a) Radiation dose rate as measured by the integrated Geiger counter and cumulative radiation dose for the duration of spaceflight. (b) Recorded temperature and humidity for the duration of spaceflight. (c) Radiation dose rate varies according to the (d) Orbital position of the ISS. (e) Images of GC and (f) FL Petri dish cultures post-flight.

Upon visual observation in comparison to ground controls, all solid cultures that were exposed to spaceflight conditions showed morphological differences, specifically in colony shape and pigment distribution (Fig. 2e, Fig. 2f). These images were captured directly after the experimental period upon opening the payload, prior to cold storage of both solid and liquid samples. Each of the post-flight experiments uses samples revived from cold storage on the dates indicated in Figure 1a and uses either the original sample (R^0^) or a single subsequent generation of regrowth (R^1^) for analysis.

### Growth studies of B. subtilis liquid samples

Growth studies measuring optical density (OD) at 600nm were conducted in order to examine the impacts of pigment shields on the R^0^ generation from liquid samples of *B. subtilis* exposed to the spaceflight environment on the ISS (FL) against matched ground control samples (GC). All studies were conducted in duplicate for each sample. Data from the growth assay were analyzed with respect to a specific set of parameters: the maximum OD, the maximum growth rate (MGR), the time to attain MGR, the time to stationary phase, and the time between MGR and stationary phase (Fig. 3a). The data were analyzed in different groupings to account for the various types of potential shields and their associated controls: pigmented (e.g., *A*.*niger*) and non-pigmented (e.g., *B. subtilis*) living microbial cultures, chemical pigment (e.g., lycopene) and non-pigment (e.g., L-tyrosine) chemical solutions, and melanin or carotenoid-containing solutions and those without pigment. The various controls allow for the examination of different shielding layers, such as the vial container, solvent, chemical pigments, and living organisms, within both the experimental and ground control payloads. The effects of spaceflight can be interpreted as a comparison of the delta value between each FL sample to its corresponding GC control sample, as variability was present across samples.

**Figure 3.**
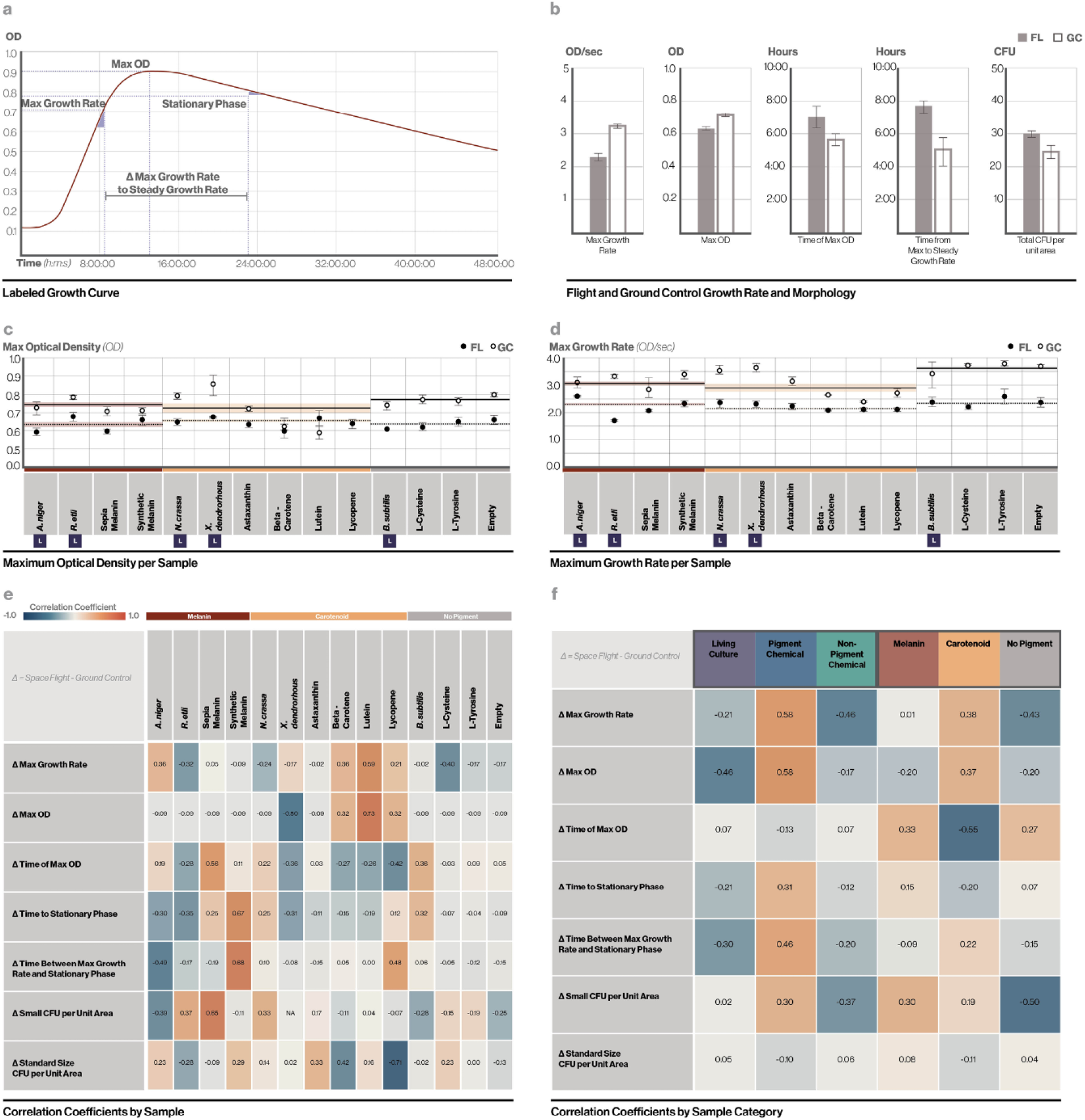
Liquid sample analysis. (a) Labeled growth curve with typical measurements of maximum OD, maximum growth rate (MGR), time to MGR,, and the time between the MGR and stationary phase. (b) B. subtilis samples from inner vials exhibited a significant difference after FL and GC conditions for the metrics of: MGR, maximum OD, time to MGR, time from MGR to stationary phase, and total CFU per unit area. Error bars represent standard error of the mean. (c) Spaceflight induced changes in max OD and (d) MGR varied according to outer vial contents. Error bars represent standard error of the mean. Horizontal lines represent the mean values across pigment groupings Melain, Carotenoid, and No Pigment, for FL (solid line) and GC (dashed line) samples, and the shaded region superimposed on each line represents the standard mean error per pigment grouping (light brown, yellow, and gray, respectively). Below the x-axis, the “L” symbol indicates pigments composed of living cultures. (e) Correlation coefficients were calculated for each sample as related to changes in growth kinetics and morphology as an indication of potential shielding effects. (f) Correlations were calculated based on sample groupings, indicating that pigment chemicals and specifically carotenoids in outer vials were correlated with shielding effects.

Both spaceflight and the presence of certain pigment materials resulted in significant differences in growth rate. Spaceflight decreased the maximum OD and MGR of *B. subtilis* across all samples and increased the times to reach the MGR and stationary phase (in a two-sample t-test, p<0.05 for each metric) (Fig. 3b). The delta between matched FL and GC samples across parameters was identified as a key measure of environmental stress mitigation by shields, with a smaller delta suggesting a greater shielding effect. As shown in Fig. 3c-d, the deltas in both MGR and maximum OD were decreased by melanin and carotenoid pigment shields, suggesting that both were effective at minimizing impacts from environmental stress. However, in both time parameters (Fig. S2), only carotenoids showed a decrease in the delta, suggesting that the two pigment types yield different effects. The melanin and carotenoid pigment categories comprise various living and non-living materials; Fig. 3c-d present values for each specific shield. Three carotenoids, beta-carotene, lutein, and lycopene, appear to have the greatest shield effects, with the smallest deltas between FL and GC samples. The delta between the FL and GC was consistent for all the non-pigment controls.

The heatmaps in Fig. 3e and 3f serve as a contextual tool to examine the relationship between shield types and the growth parameters with respect to the key measure of the delta between FL and GC samples. The color-coded values indicate if the delta is higher or lower than the mean delta across all conditions; a correlation coefficient of 1 or −1 indicates a larger comparative difference in the delta in a positive or negative direction, respectively. Pigment chemicals appear to have the strongest correlations across most parameters, suggesting that this category has the highest potential as shields. In particular, the heatmap emphasizes the reduction in the mean spaceflight-induced changes to maximum optical density and growth rate through shielding with carotenoids.

### Morphology studies of liquid samples

A count of colony-forming units (CFUs) was conducted on the R^0^ generation as a gold standard metric to compare viability between FL and GC samples. All morphology studies were done in triplicate for each sample. Three distinct colony morphologies were identified in samples sourced from the FL ancestor strain R^0^, shown in Fig. 4b: small colonies (<50% of standard size), standard sized colonies (∼1-mm diameter), and large colonies (>200% standard size). A total of five combinations of these morphologies were found across plates, and these phenotypes were conserved in the R^1^ generation, indicating a genome-level change associated with spaceflight exposure resulting in a heritable morphotype (Fig. 4a). The small colony variant (SCV) morphotype may be a marker for environmental stress experienced in spaceflight, as it was found almost exclusively on FL samples. SCVs were most consistently found in the samples shielded by chemical pigment solutions, suggesting that these may shield from one environmental stressor but not from others, triggering the expression of this morphotype (Fig. 4c). Additionally, SCVs were found in at least one duplicate of every shield type flown, except for the *A. niger* live culture, indicating its occurrence as a diversification strategy. However, small colonies may also be due to slower growth rates for FL samples (Fig. 3b, d, Supp. Fig. 2).

**Figure 4.**
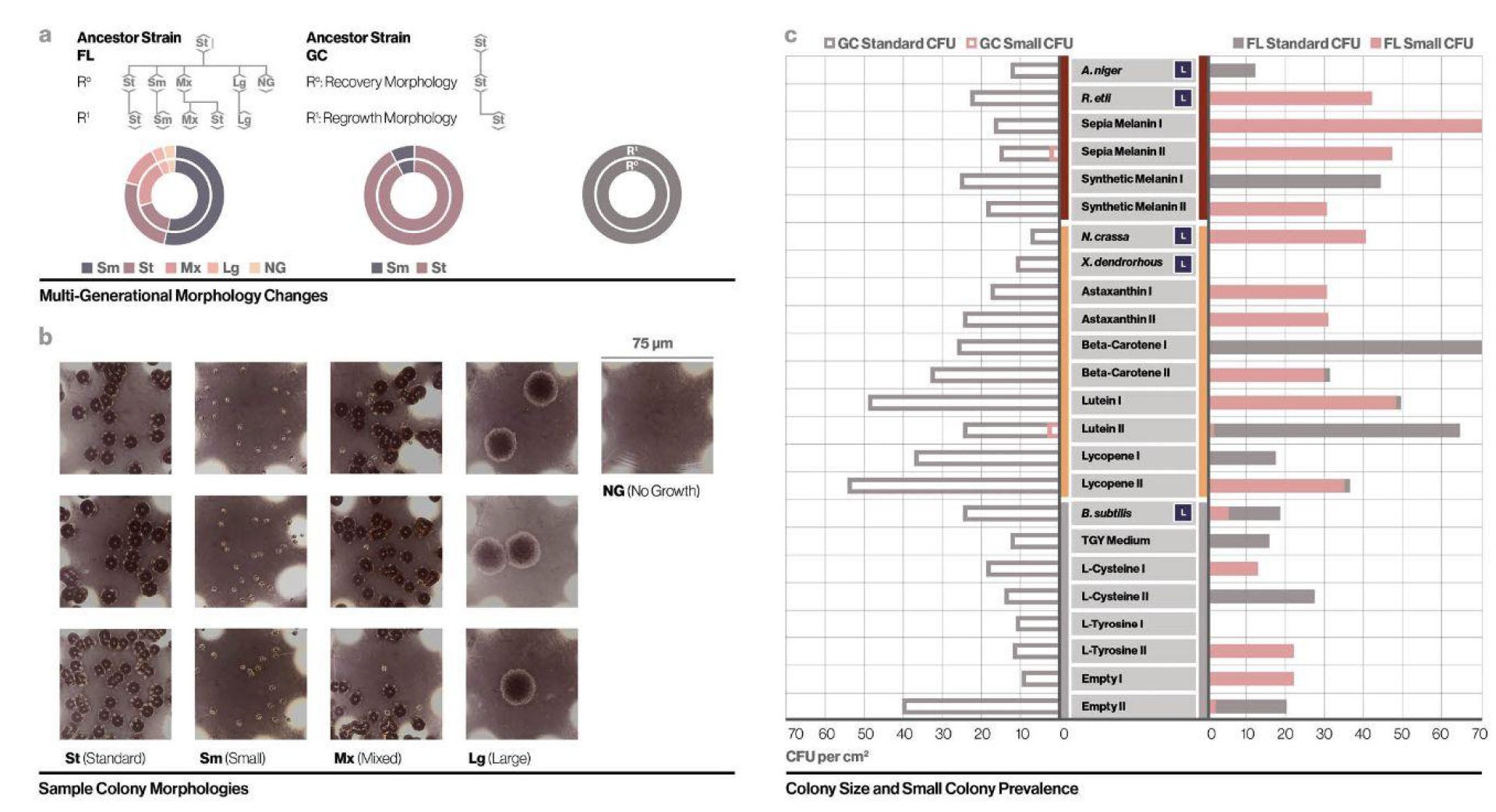
Analysis of liquid culture colony morphology. (a) Micrographs of multi-generational morphology changes in regrowth plates of B. subtilis FL liquid cultures, exhibiting three distinct colony sizes. (b) Sample morphologies over successive generations (R^0^, R^1^) for FL and GC samples (originating from the same ancestral strain), and relative proportion of each colony morphology among first and second generation. (c) Colony count and morphology make-up of each individual sample show that populations are usually skewed to one dominant morphology on a single plate.

### Growth and morphology studies for solid samples

Radial growth assays were conducted in triplicate on solid media to evaluate the growth rate, pigmentation, and morphology of the solid samples of *B. subtilis, N. crassa*, and *A. niger*. An R^1^ generation sourced from each ancestor strain was grown on fresh media and imaged once daily for eight days. Colony area, color, and shape were subsequently analyzed using Fiji^46^. Spaceflight samples of all three species were found to have a higher growth rate and colony area (Fig. 5a-c). In the case of *B. subtilis*, this is a notable difference to the liquid samples, in which spaceflight was correlated with a decreased growth rate and final concentration (Fig. 3b, d). The distinctly different biophysical environments impact how *B. subtilis*, a motile species, can move as well as access nutrients. The results for the fungal species align with previous reports of enhanced growth rates in *Penicillium* and *Aspergillus* after exposure to spaceflight in comparison to terrestrial strains^31^, and provide the first such proof for *N. crassa*, a model fungus that has rarely been studied aboard the ISS or in certain space-relevant conditions such as microgravity. The behaviors of both filamentous fungi examined in the present study support the idea that increased growth rate is an adaptation that may confer a selective advantage in stressful environments.

**Figure 5.**
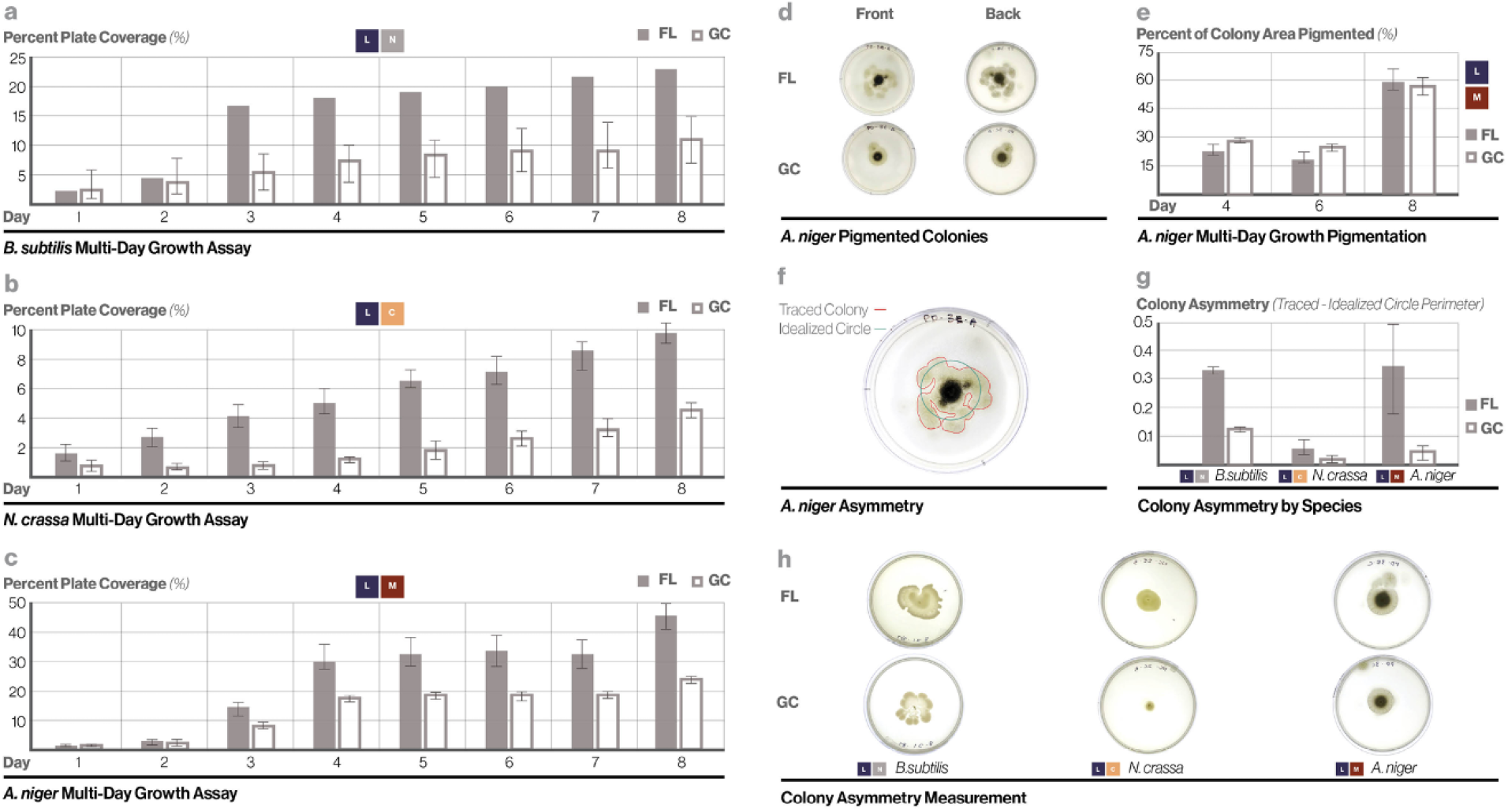
Analysis of solid culture colony morphology. (a-c) Radial growth assays of solid culture regrowths show increased colony area in each species: B. subtilis, A. niger, and N. crassa. Error bars represent standard error of the mean. (d-e) Photographic and quantitative comparison of pigmented areas of colony in FL and GC regrowths of A. niger showed minimal differences across the multi-day growth assay. Error bars represent standard error of the mean. (f-h) Photographic and quantitative comparison of colony asymmetry and irregularity in FL and GC regrowths in all three species shows significantly higher asymmetry in FL samples. Error bars represent standard error of the mean.

While slight differences in pigment distribution in *A. niger* samples were present in the ancestor strain and R^0^ generation, the R^1^ generation used in the multi-day growth assay displayed little difference between FL and GC samples (Day 4 p value = 0.203; Day 6 p value = 0.166; Day 8 p value = 0.666), indicating that this change was not conserved across generations (Fig. 5d-e). Significant changes in colony shape, most notably in symmetry, were also visually observed during multi-day radial growth assay (Fig. 5h). Qualitative analysis of *A. niger* and *B. subtilis* indicated that FL samples had visibly higher radial asymmetry and colony contour irregularity compared to controls, indicating that these features may serve as a key biomarker for spaceflight-induced stress in these species and potentially more broadly. These are measurable phenotypes that are clearly distinguishable from the radial symmetry characteristic of many microorganisms, and asymmetry has previously been identified as an indicator of environmental stress^47^. In order to quantify these morphological changes, each colony was traced in Fiji (Fig. 5f) to measure area (*A*), perimeter (*P*), and convex perimeter (*P*_*c*_). The relative difference between the perimeter of the measured colony (*P*) and the perimeter of an idealized circle of equivalent area (*P*_*i*_) was calculated as a measure of radial asymmetry (Fig. 5g, Eq. 1 (see methods)). This measurement differs from previously published methods for colony asymmetry quantification^48^ as it does not require identification of a center or orthogonal axes, potentially offering greater objectivity in measurement. In addition, roundness (Eq. 2) and convexity (Eq. 3), two metrics commonly used in shape analysis^49^, were calculated for FL and GC samples. The results indicated a significant difference in roundness for *A. niger* colonies (p = 0.019) and an insignificant difference for *N. crassa* colonies (p = 0.387); not enough *B. subtilis* FL samples were available for statistical analysis of roundness (FL = 0.527; GC average = 0.924). There was an insignificant difference in convexity between FL and GC samples for *A. niger* (p = 0.243) and *N. crassa* (p = 0.687); not enough *B. subtilis* FL samples were available for statistical analysis of convexity (FL = 0.781; GC average = 0.931).

## Discussion

In this study, we created a custom autonomous payload for the support of liquid and solid microbial samples aboard the ISS. We analyzed phenotypic changes of a common microbial model organism, with and without shielding by pigment solutions, after 30 days in spaceflight. The results indicate that there are several spaceflight-associated phenotypic changes of *B. subtilis* grown in liquid culture, including growth rate, maximum cell density, and colony size, and some of these changes are minimized by the application of pigment solution shields—most notably chemical carotenoid solutions. Environmental stress may be mitigated by solutions of both carotenoids and melanins that have been shown to effectively serve as protective shields for *B. subtilis*. In contrast to existing research on pigmented microorganisms in space and space-relevant conditions, this study provides novel proof that pigments can be applied externally as a material, rather than only produced internally by cells, to minimize the effects of environmental stress. Furthermore, it expands the nascent field of pigments as shields for space protection to include carotenoids in addition to melanins, providing a foundation for future research relevant to manned missions. We identified for the first time small colony and large colony variants in *B. subtilis* associated with spaceflight, suggesting that diversification is a strategy for long-term survival in this complex environment. We also identified colony irregularity as a key measure of spaceflight-associated phenotypic change that is present in *B. subtilis* and *A. niger* grown in solid culture. Together, these findings suggest that these microorganisms employ specific strategies, measurable through the presentation of distinct morphotypes, to overcome the challenges of spaceflight.

Here, we discuss interpretations of the results in the context of relevant literature. The history of microbial research in spaceflight environments has illustrated that the type of microorganism, its state, culture conditions, treatment across the mission, and containment strategy all affect results of phenotypic assays^26^. These complexities have made the direct interpretation of the effect of spaceflight on microorganisms difficult. In the case of *B. subtilis*, a majority of published studies have focused on spores, which demonstrate a remarkable survival response under harsh space conditions^50–52^. In this study, we chose to utilize *B. subtilis* cells in liquid media in order to explore diverse responses to environmental stress and potentially higher nutrient availability^28^. Our analysis of key phenotypic measures, most notably MGR and maximum OD, indicate that spaceflight has a significant effect on liquid cultures of *B. subtilis* (Fig. 3b). This finding, placed in the context of previous studies on *B. subtilis* in LEO in which growth rates of cells grown from spores were similar to ground controls^26^, and of those demonstrating decreased growth rates in *B. cereus* cultured in semi-solid media in spaceflight^53^, suggest that the culture condition plays a significant role in resilience to environmental stress.

Our results demonstrate pronounced differences in the shielding capabilities of chemical solutions versus biological solutions, as well as melanins versus carotenoids. Three of the carotenoid pigment solutions (beta-carotene, lutein, and lycopene) showed a significant decrease in the delta between maximum OD and MGR of spaceflight and ground control samples compared to in-payload controls; they showed consistent effects with respect to one another. This decrease suggests that basic replication mechanisms are functioning more normally when shielded by carotenoid chemical pigments; this is also supported by the maximum growth rate results (Fig. 3d). The melanins, while all demonstrating decreases in delta maximum OD compared to the in-payload controls, have a less pronounced and less consistent shielding effect. The living culture of *A. niger* appeared to cause a decrease in the delta between FL and GC for measured MGR (Fig. 3d), a finding relevant to recent studies of other melanized fungi flown on the ISS in the context of radiation shielding^17^. However, as only one sample of *A. niger* was included in this experiment and a different method of measuring shield effectiveness was utilized, more thorough studies would be required to validate this result. Pigment chemicals as a category demonstrated greater shielding effects than living samples or non-pigment chemicals. The difference between living and non-living shields may be due to the higher concentration of pigments in non-living solutions, as well as the rate of pigment production in some living samples, such as *N. crassa*, when subject to light exposure^54^ and temperature^55^. While we assume that a minimization of growth rate reduction relative to GC represents an effective shielding from environmental factors in space, slowed metabolisms are known survival strategies that may also be aided by pigments.

Morphology studies show that survival adaptations may affect growth dynamics in various ways; for instance, small colonies may correlate to lower growth rates, but can represent a strategy for increased survival that may or may not be influenced by pigment. Multi-generational phenotypic changes, particularly the emergence of small colony variant (SCV) morphologies, were present in nearly all FL samples and in almost no GC samples (Fig. 4a). This finding aligns with the occurrence of the SCV phenotype previously described in various bacteria, including *B. subtilis*, which has been associated with a variety of metabolic alterations that lead to slow growth rates, atypical biochemical characteristics, altered pathogenicity, and abnormal growth requirements ^56^. Bacterial SCVs typically form a subpopulation after exposure to environmental stress and are recognized as a distinct set of morphotypes that may vary based on the type of environmental stimulus from which they result^57^. Other species of *Bacillus* have been found to switch into SCV state after exposure to antibiotics, indicating that this morphotype may be a diversification strategy that serves as an alternative to sporulation^58^. Nearly all shielded samples had SCVs present in one or both duplicates (Fig. 4c). The exceptions were in two living pigmented shielded samples that did not have duplicates, *X. dendrorhous* and *A. niger*. This indicates that the environment of spaceflight triggered the SCV state across shield conditions, and the presence of SCVs across generations suggest that it is a heritable trait that becomes stable after initial occurrence. Our findings align with previous reports of SCVs as a form of phenotypic switching in bacteria, which is a heritable “bet-hedging” strategy and stochastic phenomenon^59^. To our knowledge, this is the first study in which SCVs are reported in spaceflown microorganisms and adds to the recent reports that catalogue microbial phenotype switching in response to spaceflight stressors^60^.

There has been increasing interest in the production of secondary metabolites, including pigments, by microorganisms in spaceflight. Several recent studies have examined members of the *Aspergillus* genus, finding changes DHN-melanin and pyranonigrin in *A. niger* isolated from the ISS^31,61^, as well as increased production of anthraquinone pigment in *A. nidulans* flown for 4-7 days aboard the ISS^62^, behaviors that have been proposed to protect from oxidative stress and radiation. Here, we found little difference in visible pigmentation in solid cultures of microorganisms studied in the radial growth assay. This finding indicates that pigment production, and more broadly, secondary metabolite production, may not be visibly impacted by short-duration spaceflight and more sensitive detection methods are required to evaluate changes.

We applied various methods to analyze and quantify morphological changes between spaceflown and ground control microbial samples (Fig. 5), and we identified asymmetry and irregular contours of colonies as potential markers of spaceflight-induced stress. These morphologies represent a distinct deviation from “standard” growth patterns and have previously been associated with various environmental stressors, such as high solar radiation^30,48,63^, and are consistent with recent studies of stress-induced morphological plasticity in bacteria^64^. We found that radial asymmetry and irregularity were clearly observable in spaceflown colonies of *A. niger* and *B. subtilis*, yet not observed in *N. crassa*. While further studies are required to unravel the mechanism behind these morphologies and their occurrence in some microorganisms and not others, they are easy to observe and quantify in setups such as those presented here and are therefore relevant to future microbiological studies in space.

While the autonomous payload functioned effectively throughout the experimental period, we propose several improvements to ensure robust image capture and environmental control. The M4 camera lenses used for active imaging were focused by screwing them until the lens was the correct distance from the sensor to provide a clear image. This configuration did not adequately secure the lens during the launch conditions, and the resulting time-lapse images were unfocused. This issue could be addressed by the use of an auto-focusing camera module, fixing the lens position more securely, or by increasing the imaging system’s depth-of-field. The active imaging system was further impaired by the development of condensation in the sealed culture plates. This can be attributed in part due to the higher-than-expected temperatures within the capsule. Integrated temperature and humidity sensors recorded temperatures nearing 40°C and over 90% humidity within the capsule. The elevated temperature may have resulted from the operations of neighboring payloads, as several of these employed active lighting and on-board computers, both of which could generate substantial heat throughout the experiment. Such a transfer of heat is a common issue with similar payloads^65,66^. The most significant source of this heat, however, was the capsule’s integrated microcomputers, which could be mitigated with the use of heat sinks to potentially maintain an ideal microbial growth temperature (∼30°C) in future deployments. The Raspberry Pi microcomputers within our capsule were programmed to operate in an idle mode when not collecting ambient data or taking photos, which occurred at five-minute intervals. This configuration was programmed to reduce heat generation within the capsule, but temperatures of over 30°C were still recorded during tests in ambient conditions over recording periods of several days. Creating an effective temperature control is a distinct challenge, given that launch conditions required the capsule to be water-tight and samples at least double contained. Conventional cooling systems such as those using fan arrays cannot be applied to the enclosed system. An external water circulation system, such as those used in high-end personal computers, could be used to maintain a constant temperature. However, such a system would further reduce the available space for other components and experiments within the capsule.

Automated, inexpensive payload platforms that can collect data without astronaut interaction hold remarkable potential to advance microbiological research in spaceflight environments, especially considering the increasing accessibility of the ISS to experimenters. Within these payload systems, visible phenotypes such as colony macro-morphologies can be feasibly measured and quantified, enabling real-time analysis of the impact of spaceflight and mitigating strategies. A catalogue of microbial phenotypes and key markers of environmental stress can provide a crucial baseline for future experiments of this kind. We propose that growth rate, maximum OD, the presence of small colonies, and asymmetry may all serve as indicators of spaceflight induced stress.

This is the first study to investigate the use of liquid pigment solutions as potential shielding from environmental stressors aboard the ISS, and the first to utilize analysis of phenotypes to verify the utility and function of pigments both within and outside of cells. Our results show that pigment solutions minimize the impact of an array of spaceflight induced stresses, and that different pigments such as carotenoids and melanins have distinct and separate mitigating effects. These findings indicate that pigment solutions hold promise as biocompatible shields that may be complementary to recent advances in wearable protection for manned spaceflight, such as water-filled garments^67,68^. In future manned missions, pigment solutions may hold the key to protecting astronauts against the harsh environment of space.

## Materials and Methods

### Cell selection and culture

*Bacillus subtilis subtilis* (Ehrenberg) Cohn (ATCC 6051), *Aspergillus niger* van Tieghem (ATCC 16888), *Rhizobium etli* Segovia et al. (ATCC 51251), *Xanthophyllomyces dendrorhous* Golubev (ATCC 24230), and *Neurospora crassa* Shear et Dodge (ATCC 10815) were purchased from the American Type Culture Collection (ATCC). *B. subtilis* 6051 was selected as it is a common model strain and has previously been used in spaceflight experiments^41,69^. *A. niger* was selected due to previous reports that this species exists as part of the ISS microbiome^31^; strain 16888 was selected due to its melanin production and use as a common and available model. *R. etli* 51251 was selected as another melanin-producing bacteria in contrast to the fungus *A. niger*; it is a common model organism. *X. dendrorhous* 24230 was selected for its high carotenoid production. *N. crassa* was selected as it has been used previously in spaceflight research^70,71^; strain 10815 was selected due to its production of carotenoids and availability. All strains can be easily distinguished through microscopic inspection and macroscopic evaluation of color and colony morphology. All liquid cultures of *B. subtilis* were in Luria Broth medium before, during, and after spaceflight. Vial cultures of *X. dendrorhous* were in Yeast Malt Medium; *R. etli* was cultured in Rhizobium medium (ATCC); *A. niger* in Potato Dextrose Medium. Solid samples in spaceflight were cultured on LB Agar (*B. subtilis* and co-cultures), Neurospora crassa Agar (*N. crassa*), and Potato Dextrose Agar (*A. niger*). All samples were stored in 10%–50% glycerol at −80°C as soon as possible post-flight. For post-flight assays, *B. subtilis* liquid samples were cultured in LB medium at 37°C, 200rpm; solid cultures were grown on LB Agar (*B. subtilis*), Neurospora crassa Agar (*N. crassa*), and Potato Dextrose Agar (*A. niger*) at 25°C.

### Payload Design and Fabrication

The payload was designed with an optimal packing strategy to house as many vials and 50mm culture plates as possible while allowing for time-lapse imaging of the dishes. It was one of several similar-sized units within the NanoRacks BlackBox, an autonomous hardware platform for small-scale experimentation aboard the ISS. The BlackBox provided 5V power through a USB connection and an additional layer of housing and vibration protection. A novel workflow was developed that integrates a series of genetic algorithms, computational design software, and 3D printing to easily optimize the arrangement, containment, and fabrication process. Contents were modeled in the CAD software Rhinoceros 3D and were tagged as follows:

1. Fixed Components: Objects that are required to be in a specific location and will not be moved in tested configurations. The system’s USB power inlet was placed as a fixed component based on a planned symmetrical design so that the power source would be equidistant from the two powered components. Exterior chassis components were defined as fixed components due to their predetermined configuration, given by the requirements of the NanoRacks BlackBox.
2. Grouped Components: Objects that can be rearranged so long as their orientation relative to one another remains constant. These included the camera sensor, lens, and culture plates. These components were grouped so that regardless of the group’s general position and orientation, the three culture plates would always be within the camera’s field of view. This category also proved useful for components that could be manually configured into a packed grouping before placement. In this capsule, vials containing passive experiments were manually grouped into a packed configuration with their lids remaining accessible.
3. Free-Floating Components: Objects that can be located anywhere within the arrangement. This category included sensors for temperature, humidity, and radiation. These components could feasibly be placed anywhere within the capsule without undue impact on the experimental conditions.
4. Void Spaces: Regions that must remain empty. This included the space, the camera, and culture plates to maintain a clear view of the experiments. Additional void space was designated in the capsule’s center to allow for easier access and placement of the system’s components during construction.
5. Flexible Connections: This category included wired connections between the microcomputers and sensors as well as the power input to the microcomputers. While these connections can be adjusted, they were minimized to eliminate excess lengths of wire that could obscure images or prove challenging to set into place.
6. Experimental Bounds: The space within which all contents must fit. As this payload was one of five within the BlackBox, the exterior dimensions were constrained to 2U (4 × 4 × 8 in.).

Once experimental contents were categorized, the sub-arrangement of coupled components was locally optimized. We implemented a genetic algorithm in which the camera, lens, and monitored culture plates are allowed to rotate in three planes until a configuration that minimizes the volume of an oriented bounding box was found. During this rotation, the culture-dish angle and distance of the camera are kept constant. The minimization of the bounding box ensured that the group components were occupying as little space as possible without compromising the camera’s view of the culture plates. Void space, grouped components, and free-floating components were continually rotated and translated within the experimental bounds by a second genetic algorithm. The algorithm used the Cartesian coordinates and rotation of each component in its genome and evaluated the minimal bounding box volume of all components as a fitness parameter. Configurations that intersected with the experimental bounds had their fitness penalized to encourage a configuration that fits within the bounds. When an optimal configuration had been established, objects were suspended in a configuration and required support. To fix components into place, an agent-based model was used to join components to each other and the capsule chassis while avoiding designated void spaces. The agent-based model used a modification of the Boids algorithm^72^ for simulation of flocking behavior. The paths of each agent were connected to form smoothed polylines. These polylines were then converted into volumes through the use of a signed distance function and were subsequently meshed to create objects suitable for 3D printing. A Markforged Onyx One desktop 3D printer was used to print the capsule from filament containing nylon and chopped carbon fiber (Onyx). This material was selected for its durability and flexibility, allowing for the box to be slightly deformed without causing damage. Electrical components were then friction fit into the printed capsule or were secured with a clear silicone sealant (Loctite).

### Payload Preparation

The payload was leak tested following Methods VIII and IX from Lvovsky & Grayson (ARES Corporation). Biological samples were included in this experiment in two formats: as liquid culture in nesting polypropylene vials (VWR) and as solid agar cultures in 50mm polystyrene dishes (Thermofisher Scientific). The nesting vials comprised a 2-mL screw-top vial, which contained liquid *B. subtilis* culture (“inner”), placed inside a 5-mL screw-top vial (“outer”), which contained the pigment solutions and associated controls. *B. subtilis* cultures were grown overnight in LB broth at 150rpm, 30°C; and 2-mL was filled into each of the inner vials. 3-mL of each pigment solution (1g/1-mL) was filled into each outer vial. The solid agar plates contained 12-mL of 2% agar with corresponding nutrients (LB Agar for *B. subtilis* and co-cultures, Neurospora Crassa Agar (ATCC) for *N. crassa*, Potato Dextrose Agar for *A. niger*). *B. subtilis, N. crassa*, and *A. niger* colonies were picked from plates stored at 4°C and seeded in the center of each solid culture plate. Plates were closed and sealed with a single layer of Parafilm (VWR) to allow for water containment and gas permeability. One hour before shipping, all vials and dishes were fit into the payload and secured as needed with silicone adhesive (Loctite). The adhesive was also applied to the lens rings of the cameras to fix the focus. The payload was closed and polyimide tape (Kapton) was applied to the seam to ensure a proper layer of containment. An elastic band was further applied to slightly compress the payload.

### Mission timeline and profile

Samples were inoculated and loaded on February 21, 2020, and the payload was shipped from Cambridge, MA to Houston, TX. The payload was integrated into the NanoRacks BlackBox on February 22. Due to launch rescheduling, they remained contained with the payload unpowered at ambient conditions until March 6. The SpaceX CRS-20 Dragon spacecraft launched aboard a Falcon 9 rocket from Cape Canaveral, FL on March 6 and arrived at the ISS on March 9; the BlackBox was plugged into power shortly thereafter. During spaceflight, the temperature range was 24.5°C - 41.6°C and the humidity range was 32.10% - 99.60%, as recorded by the in-payload sensors in five-minute intervals. After a 30 day experimental period, the Dragon departed the ISS on April 7 and splashed down west of Baja California in the Pacific Ocean that morning. The BlackBox was opened on April 13 in Houston, TX and the payload was received in Cambridge, MA on April 14. All biological and chemical samples were stored in 10-50% glycerol and placed in −80°C on April 17. Analysis began in June 2020 due to delays from the Covid-19 pandemic.

### Vials: Recovery and OD_600_ growth assay

Inner vial samples of *B. subtilis* (V1-24, Experimental and Control) were recovered on L+91 from stocks stored at 4°C. Samples were diluted 1:10 in Nutrient Broth and dispensed in duplicate (n=2, 200-μl each) in a glass-bottomed 96-well plate. Two blank wells comprising Nutritional Broth were included. A BioTek Synergy 2 plate reader provided shaking incubation at 30°C and recorded OD_600_ absorbance measurements at 10-minute intervals for 48 hours. To produce growth curves, we passed the OD_600_ time-course data per well through a 10-point rectangular smoothing function and subtracted the smoothed, averaged blank well OD_600_ values per time point. The following metrics, used to characterize growth dynamics, were derived via the growth curves using Microsoft Excel: Peak Culture Density (i.e., maximum OD), Maximum Growth Rate (MGR), Time to MGR, and Time to Stationary Phase.

The Peak Culture Density was the maximum OD value for each well’s growth curve. For “MGR”, we identified the maximum positive value (unit: ΔOD/sec) of the first derivative of each well’s growth curve (i.e., OD’ with respect to time). For the “Time to MGR” metric, the start of incubation was defined as t=0 and a time value (unit: hours) corresponding to the maxima of the OD’ curve was identified. Finally, the first time point at which the OD’ = 0 was identified; the “Time to Stationary Phase” represents the value (unit: hours) from t=0 until this time point, while “Time from MGR to Stationary Phase” metric represents the value (unit: hours) from the curve’s corresponding time of maximum growth rate until this time point. For each of these growth metrics, values were further grouped by vial type, creating a mean and variance value (i.e., standard mean error) for each metric per shielding material (n=4 for “Non-living” vial types, n=2 for “Living” vial types).

### Vials: Colony Counts and Microscopy, Quantitative Analysis

Vial samples of *B. subtilis* (V1-24, E, and C) were recovered on L+186d from glycerol stocks stored at −80°C and streaked onto plates of Nutrient Agar, 2% (BD Biosciences). Recovery plates (R^0^) incubated for 23 hours at 30°C. Morphologies observed on recovery plates were noted and each plate was imaged by a DSLR digital camera (Canon Mark IV). Single colonies from each recovery plate were picked and used to inoculate a 2ml liquid culture of Nutrient Broth (BD Biosciences). Liquid regrowth cultures (R^1^) incubated for 16 hours at 30°C and 200 rpm. Liquid cultures were then diluted 1:10^6^ in Nutrient Broth and plated in triplicate, with 100 mL of the diluted culture evenly spread with a disposable sterile L-shaped spreader on each plate (Nutrient Agar, 2%). Dilution plates incubated for 16 hours at 30°C and then were imaged for colony count and morphology analysis.

For automated (macro) colony counting, plates were imaged by a DSLR digital camera (Canon Mark IV) and counted in Fiji with the Analyze Particles tool, after manual thresholding and ROI selection. For manual (micro) colony counting and verification, four random magnified selections (10 × 7.5 mm) of each plate were imaged by a USB digital microscope (Plugable), and colonies of the distinct phenotype were manually classified and counted. A colony count per area value was averaged across the four image samples per plate.

### Plates: recovery and multi-day growth assay, quantitative and qualitative Analysis

Solid plate culture samples (P1-6, E, and C), comprising *B. subtilis, N. crassa*, and *A. niger* in mono- and co-culture arrangements, were recovered on L+202d from glycerol stocks stored at −80°C. For recovery (R^0^), stocks were thawed momentarily via 37°C water bath and streaked onto plates of respective media per species; co-culture plates used Nutrient Agar. Recovery plates incubated at 25°C for 5 days. Morphologies observed on recovery plates were noted and photographically recorded with a Canon Mark IV DSLR Camera. For regrowth (R^1^), a stab from each recovery plate was taken via a sterile pipette tip and transferred to the center of a regrowth plate of the same media. Regrowth plates incubated for 8 days at 25°C. Plates were imaged from a bottom-view every 24 hours (Days 0-8); plates were also imaged from directly above on Days 4, 6, and 8.

Central colonies of both monoculture and co-culture plates were manually outlined in Fiji in order to measure colony area in pixels and plate area in pixels. Percent cover ((Colony Area/Plate Area) x 100) was calculated for each plate; averages and standard error of the mean were computed for replicate samples. Images of plates that had excess condensation leading to disruption of colony formation were excluded from all analyses. All experts conducting manual outlining had 95%+ agreement. Grids were created for qualitative analysis of spaceflown plate samples, recovery (R^0^) plates, and multi-day assay monoculture and co-culture (R^1^) plates.

Images of the front of monoculture plates obtained on Day 3 of the multi-day growth assay were utilized for symmetry analysis. Central colonies of these plates were manually outlined in Fiji in order to measure colony perimeter, convex perimeter (using the *convex hull* function), and area. Three metrics were utilized to quantify asymmetry and irregularity: asymmetry (Eq. 1), roundness (Eq. 2), and convexity (Eq. 3).

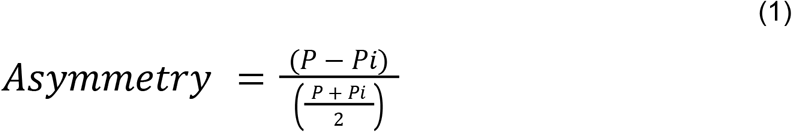

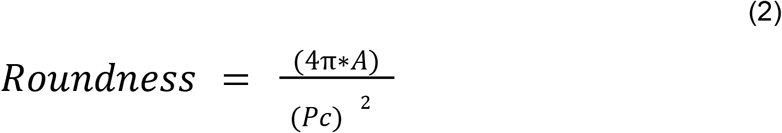

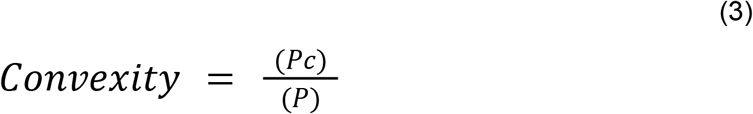

Images of the front of *A. niger* monoculture plates obtained on Day 3, Day 5, and Day 7 of the multi-day growth assay were utilized for visible pigmentation analysis. Central colonies of these plates were manually outlined in Fiji in order to measure colony area; the pigmented area, appearing black, was also manually outlined and area was measured. Percent pigmented ((Pigmented Area/Colony Area) x 100) was calculated for each plate; averages and standard error of the mean were computed for replicate samples.

### Statistical analysis

Statistical analysis and graph generation were accomplished using Sheets (Google), Excel (Microsoft), and Illustrator (Adobe). For analyses comparing FL and GC conditions, two-tailed, two-sample Welch’s unequal variances t-tests were performed with a p-value threshold of 0.05. For analysis comparing melanin, carotenoid, and non-pigment categorical groupings, single factor ANOVA tests were performed with a p-value threshold of 0.05.

## Author Contributions

S.S. conceived the study, created protocols for pre-flight and post-flight testing, integrated the payload, and received and stored the samples post-flight. S. S., S. W., R. S. H. S., and N. L. conducted pre-flight testing and prepared the payload for launch. S. S. and R. S. H. S. conducted post-flight assays. S. S., R. S. H. S., and N. L. participated in data analysis, statistical analysis, and creating figure content. N. L. formatted the final figures. S. S. wrote the initial manuscript draft; N. L., R. S. H. S, S. W., and N. O. provided editing. N. O. provided day-to-day oversight and support for this research. S. S. and N. O. were co-investigators for the NASA-TRISH-funded project “Biological Pigments for Space Radiation Protection”. S. S. and R. S. H. S. contributed equally to this work.

## Data availability

The datasets and images generated during and/or analyzed during the current study are available from the corresponding author on request.

## Acknowledgments

This work was made possible through collaboration with the MIT Space Exploration Initiative, which provided a slot for the experiment payload within the NanoRacks BlackBox on the SpaceX CRS-20 flight. Funding was provided by the MIT-SEI Translational Research in Space Health Seed Fund and the MIT Media Lab Consortium. The authors acknowledge A. E. and J. S. for helpful discussions and M. M. for assistance with payload integration.

## Ethics declarations

The authors declare they have no competing financial interests.

## References

1. Microbiology Program. https://science.nasa.gov/biological-physical/programs/space-biology/microbiology.

2. Novikova, N. et al. Survey of environmental biocontamination on board the International Space Station. Res. Microbiol. 157, 5–12 (2006).

3. Coil, D. A. et al. Growth of 48 built environment bacterial isolates on board the International Space Station (ISS). PeerJ 4, e1842 (2016).

4. Lang, J. M. et al. A microbial survey of the International Space Station (ISS). PeerJ 5, e4029 (2017).

5. Mora, M. et al. Resilient microorganisms in dust samples of the International Space Station-survival of the adaptation specialists. Microbiome 4, 65 (2016).

6. Mora, M. et al. Space Station conditions are selective but do not alter microbial characteristics relevant to human health. Nat. Commun. 10, 3990 (2019).

7. Bijlani, S., Stephens, E., Singh, N. K., Venkateswaran, K. & Wang, C. C. C. Advances in space microbiology. iScience 24, 102395 (2021).

8. Cucinotta, F. A. Radiation risk acceptability and limitations. Space Radiation Program Element, NASA Johnson Space Center, Houston, TX (2010).

9. Bagatin, M. Ionizing radiation effects in electronics. (2015) doi:10.1201/b19223.

10. Cordero, R. J. & Casadevall, A. Functions of fungal melanin beyond virulence. Fungal Biol. Rev. 31, 99–112 (2017).

11. Cordero, R. J. B., Vij, R. & Casadevall, A. Microbial melanins for radioprotection and bioremediation. Microb. Biotechnol. 10, 1186–1190 (2017).

12. Zhdanova, N. N., Zakharchenko, V. A., Vember, V. V. & Nakonechnaya, L. T. Fungi from Chernobyl: mycobiota of the inner regions of the containment structures of the damaged nuclear reactor. Mycol. Res. 104, 1421–1426 (2000).

13. Zhdanova, N. N., Tugay, T., Dighton, J., Zheltonozhsky, V. & McDermott, P. Ionizing radiation attracts soil fungi. Mycol. Res. 108, 1089–1096 (2004).

14. El-Naggar, N.E.-A. & El-Ewasy, S. M. Bioproduction, characterization, anticancer and antioxidant activities of extracellular melanin pigment produced by newly isolated microbial cell factories Streptomyces glaucescens NEAE-H. Sci. Rep. 7, 42129 (2017).

15. Martínez, L. M., Martinez, A. & Gosset, G. Production of Melanins With Recombinant Microorganisms. Front Bioeng Biotechnol 7, 285 (2019).

16. Cordero, R. J. B. Melanin for space travel radioprotection: Melanin radioprotection. Environ. Microbiol. 19, 2529–2532 (2017).

17. Shunk, G. K., Gomez, X. R. & Averesch, N. J. H. A Self-Replicating Radiation-Shield for Human Deep-Space Exploration: Radiotrophic Fungi can Attenuate Ionizing Radiation aboard the International Space Station. bioRxiv 2020.07.16.205534 (2020) doi:10.1101/2020.07.16.205534.

18. Scientists send melanin into space for six months. The Johns Hopkins News-Letter https://www.jhunewsletter.com/article/2019/11/scientists-send-melanin-into-space-for-six-months (2019).

19. Gajowik, A. & Dobrzyńska, M. M. Lycopene - antioxidant with radioprotective and anticancer properties. A review. Rocz. Panstw. Zakl. Hig. 65, 263–271 (2014).

20. Srinivasan, M. et al. Lycopene as a natural protector against γ-radiation induced DNA damage, lipid peroxidation and antioxidant status in primary culture of isolated rat hepatocytes in vitro. Biochimica et Biophysica Acta (BBA) - General Subjects 1770, 659–665 (2007).

21. Srinivasan, M., Devipriya, N., Kalpana, K. B. & Menon, V. P. Lycopene: An antioxidant and radioprotector against gamma-radiation-induced cellular damages in cultured human lymphocytes. Toxicology 262, 43–49 (2009).

22. Dartnell, L. R. et al. Destruction of Raman biosignatures by ionising radiation and the implications for life detection on Mars. Anal. Bioanal. Chem. 403, 131–144 (2012).

23. Adamec, F. et al. Photophysics of deinoxanthin, the keto-carotenoid bound to the main S-layer unit of Deinococcus radiodurans. Photochem. Photobiol. Sci. 19, 495–503 (2020).

24. DasSarma, S., DasSarma, P., Laye, V. J. & Schwieterman, E. W. Extremophilic models for astrobiology: haloarchaeal survival strategies and pigments for remote sensing. Extremophiles 24, 31–41 (2020).

25. Nickerson, C. A., Ott, C. M., Wilson, J. W., Ramamurthy, R. & Pierson, D. L. Microbial responses to microgravity and other low-shear environments. Microbiol. Mol. Biol. Rev. 68, 345–361 (2004).

26. Morrison, M. D., Fajardo-Cavazos, P. & Nicholson, W. L. Comparison of Bacillus subtilis transcriptome profiles from two separate missions to the International Space Station. npj Microgravity 5, 1 (2019).

27. Afshinnekoo, E. et al. Fundamental Biological Features of Spaceflight: Advancing the Field to Enable Deep-Space Exploration. Cell 183, 1162–1184 (2020).

28. Milojevic, T. & Weckwerth, W. Molecular Mechanisms of Microbial Survivability in Outer Space: A Systems Biology Approach. Front. Microbiol. 11, 923 (2020).

29. Horneck, G., Klaus, D. M. & Mancinelli, R. L. Space microbiology. Microbiol. Mol. Biol. Rev. 74, 121–156 (2010).

30. Singaravelan, N. et al. Adaptive melanin response of the soil fungus Aspergillus niger to UV radiation stress at ‘Evolution Canyon’, Mount Carmel, Israel. PLoS One 3, e2993 (2008).

31. Romsdahl, J. et al. Characterization of Aspergillus niger Isolated from the International Space Station. mSystems 3, (2018).

32. Gomoiu, I., Chatzitheodoridis, E., Vadrucci, S., Walther, I. & Cojoc, R. Fungal Spores Viability on the International Space Station. Orig. Life Evol. Biosph. 46, 403–418 (2016).

33. Schmidhauser, T. J., Lauter, F. R., Russo, V. E. & Yanofsky, C. Cloning, sequence, and photoregulation of al-1, a carotenoid biosynthetic gene of Neurospora crassa. Mol. Cell. Biol. 10, 5064–5070 (1990).

34. Iigusa, H., Yoshida, Y. & Hasunuma, K. Oxygen and hydrogen peroxide enhance light-induced carotenoid synthesis in Neurospora crassa. FEBS Lett. 579, 4012–4016 (2005).

35. Harding, R. W. & Turner, R. V. Photoregulation of the Carotenoid Biosynthetic Pathway in Albino and White Collar Mutants of Neurospora crassa. Plant Physiol. 68, 745–749 (1981).

36. Brehm-Stecher, B. F. & Johnson, E. A. Isolation of carotenoid hyperproducing mutants of Xanthophyllomyces dendrorhous (Phaffia rhodozyma) by flow cytometry and cell sorting. Methods Mol. Biol. 898, 207–217 (2012).

37. Kim, J.-H., Kang, S.-W., Kim, S.-W. & Chang, H.-I. High-level production of astaxanthin by Xanthophyllomyces dendrorhous mutant JH1 using statistical experimental designs. Biosci. Biotechnol. Biochem. 69, 1743–1748 (2005).

38. Sharma, R. et al. The genome of the basal agaricomycete Xanthophyllomyces dendrorhous provides insights into the organization of its acetyl-CoA derived pathways and the evolution of Agaricomycotina. BMC Genomics 16, 233 (2015).

39. Cabrera-Valladares, N. et al. Expression of the melA gene from Rhizobium etli CFN42 in Escherichia coli and characterization of the encoded tyrosinase. Enzyme Microb. Technol. 38, 772–779 (2006).

40. Michiels, J., Moris, M., Dombrecht, B., Verreth, C. & Vanderleyden, J. Differential regulation of Rhizobium etli rpoN2 gene expression during symbiosis and free-living growth. J. Bacteriol. 180, 3620–3628 (1998).

41. Kacena, M. A. et al. Bacterial growth in space flight: logistic growth curve parameters for Escherichia coli and Bacillus subtilis. Appl. Microbiol. Biotechnol. 51, 229–234 (1999).

42. Morrison, M. D. Investigating the Effects of the Human Spaceflight Environment on Bacillus subtilis. (University of Florida, 2019).

43. Fajardo-Cavazos, P. & Nicholson, W. L. Establishing Standard Protocols for Bacterial Culture in Biological Research in Canisters (BRIC) Hardware. Gravitational and Space Research 4, 58–69 (2020).

44. Nicholson, W. L. Spore-forming bacteria as model organisms for studies in astrobiology. Extremophiles as Astrobiological Models 275–290 (2020) doi:10.1002/9781119593096.ch13.

45. NASA. Space Radiation.

46. Schindelin, J. et al. Fiji: an open-source platform for biological-image analysis. Nat. Methods 9, 676–682 (2012).

47. Graham, J. H., Raz, S., Hel-Or, H. & Nevo, E. Fluctuating Asymmetry: Methods, Theory, and Applications. Symmetry 2, 466–540 (2010).

48. Raz, S. et al. Growth and Asymmetry of Soil Microfungal Colonies from ‘Evolution Canyon,’ Lower Nahal Oren, Mount Carmel, Israel. PLoS One 7, e34689 (2012).

49. Wirth, M. A. Shape Analysis and Measurement. (2004).

50. Horneck, G. et al. Resistance of Bacterial Endospores to Outer Space for Planetary Protection Purposes—Experiment PROTECT of the EXPOSE-E Mission. Astrobiology 12, 445–456 (2012).

51. Horneck, G. Responses of Bacillus subtilis spores to space environment: results from experiments in space. Orig. Life Evol. Biosph. 23, 37–52 (1993).

52. Santomartino, R. et al. No Effect of Microgravity and Simulated Mars Gravity on Final Bacterial Cell Concentrations on the International Space Station: Applications to Space Bioproduction. Front. Microbiol. 11, 579156 (2020).

53. Su, L. et al. Phenotypic, genomic, transcriptomic and proteomic changes in Bacillus cereus after a short-term space flight. Adv. Space Res. 53, 18–29 (2014).

54. Luque, E. M. et al. A relationship between carotenoid accumulation and the distribution of species of the fungus Neurospora in Spain. PLoS One 7, e33658 (2012).

55. Castrillo, M. et al. Transcriptional basis of enhanced photoinduction of carotenoid biosynthesis at low temperature in the fungus Neurospora crassa. Res. Microbiol. 169, 78–89 (2018).

56. Proctor, R. A. et al. Small colony variants: a pathogenic form of bacteria that facilitates persistent and recurrent infections. Nat. Rev. Microbiol. 4, 295–305 (2006).

57. Johns, B. E., Purdy, K. J., Tucker, N. P. & Maddocks, S. E. Phenotypic and Genotypic Characteristics of Small Colony Variants and Their Role in Chronic Infection. Microbiol Insights 8, 15–23 (2015).

58. Frenzel, E., Kranzler, M., Stark, T. D., Hofmann, T. & Ehling-Schulz, M. The Endospore-Forming Pathogen Bacillus cereus Exploits a Small Colony Variant-Based Diversification Strategy in Response to Aminoglycoside Exposure. MBio 6, e01172–15 (2015).

59. Cui, L., Neoh, H.-M., Iwamoto, A. & Hiramatsu, K. Coordinated phenotype switching with large-scale chromosome flip-flop inversion observed in bacteria. Proc. Natl. Acad. Sci. U. S. A. 109, E1647–56 (2012).

60. Searles, S. C., Woolley, C. M., Petersen, R. A., Hyman, L. E. & Nielsen-Preiss, S. M. Modeled microgravity increases filamentation, biofilm formation, phenotypic switching, and antimicrobial resistance in Candida albicans. Astrobiology 11, 825–836 (2011).

61. Romsdahl, J., Blachowicz, A., Chiang, Y.-M., Venkateswaran, K. & Wang, C. C. C. Metabolomic Analysis of Aspergillus niger Isolated From the International Space Station Reveals Enhanced Production Levels of the Antioxidant Pyranonigrin A. Front. Microbiol. 11, 931 (2020).

62. Romsdahl, J. et al. International Space Station conditions alter genomics, proteomics, and metabolomics in Aspergillus nidulans. Appl. Microbiol. Biotechnol. 103, 1363–1377 (2019).

63. Nevo, E. ‘Evolution Canyon,’ a potential microscale monitor of global warming across life. Proc. Natl. Acad. Sci. U. S. A. 109, 2960–2965 (2012).

64. Ultee, E., Ramijan, K., Dame, R. T., Briegel, A. & Claessen, D. Chapter Two - Stress-induced adaptive morphogenesis in bacteria. in Advances in Microbial Physiology (ed. Poole, R. K.) vol. 74 97–141 (Academic Press, 2019).

65. Nicholson, W. L. et al. The O/OREOS mission: first science data from the Space Environment Survivability of Living Organisms (SESLO) payload. Astrobiology 11, 951–958 (2011).

66. Nicholson, W. L. & Ricco, A. J. Nanosatellites for Biology in Space: In Situ Measurement of Bacillus subtilis Spore Germination and Growth after 6 Months in Low Earth Orbit on the O/OREOS Mission. Life 10, (2019).

67. Baiocco, G. et al. A water-filled garment to protect astronauts during interplanetary missions tested on board the ISS. Life Sci. Space Res. 18, 1–11 (2018).

68. Vuolo, M. et al. Exploring innovative radiation shielding approaches in space: A material and design study for a wearable radiation protection spacesuit. Life Sci. Space Res. 15, 69–78 (2017).

69. Leys, N. M. E. J., Hendrickx, L., De Boever, P., Baatout, S. & Mergeay, M. Space flight effects on bacterial physiology. J. Biol. Regul. Homeost. Agents 18, 193–199 (2004).

70. Sulzman, F. M., Ellman, D., Fuller, C. A., Moore-Ede, M. C. & Wassmer, G. Neurospora circadian rhythms in space: a reexamination of the endogenous-exogenous question. Science 225, 232–234 (1984).

71. Ferraro, J. S., Fuller, C. A. & Sulzman, F. M. The biological clock of Neurospora in a microgravity environment. Adv. Space Res. 9, 251–260 (1989).

72. Reynolds, C. W. Flocks, Herds, and Schools: A Distributed Behavioral Model. Comput. Graph. 21, 25–34 (1987).

