## Supplementary Information for "Exogenous pigments shield microorganisms from spaceflight-induced changes"

Sharma et al.

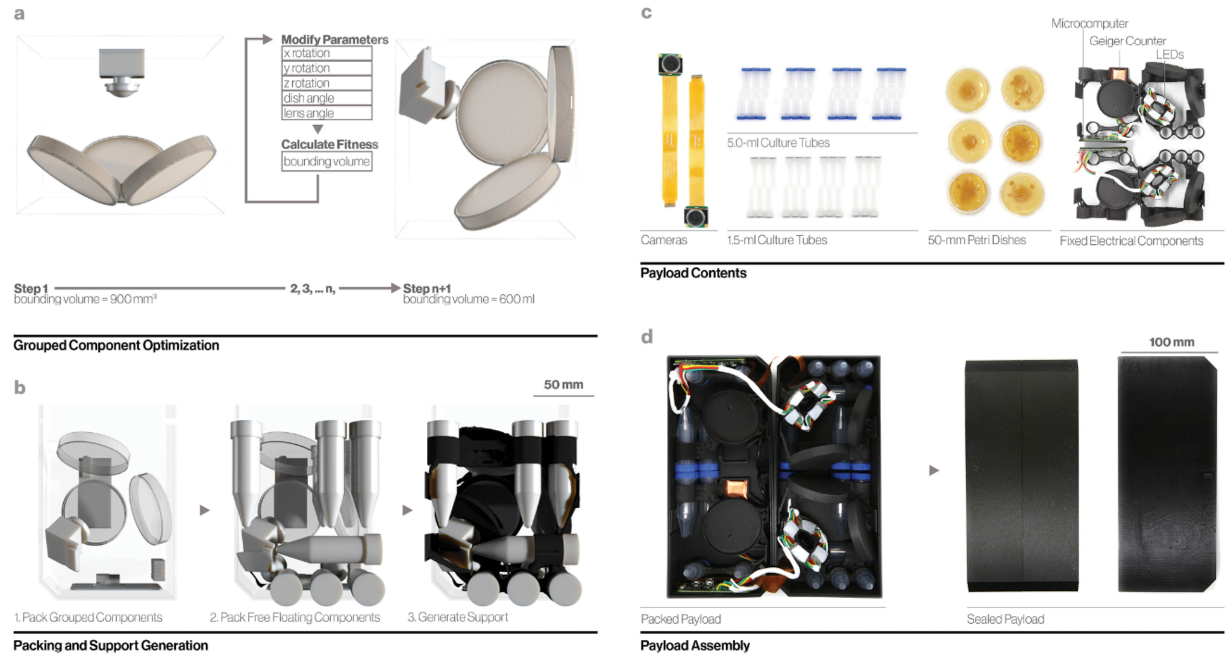


*Supplementary Figure 1. Radiofungi payload design and components. (a) Optimization of camera and Petri dish arrangement. (b) Optimized packing strategy development and support material placement. (c) Contents within the payload. (d) Final payload assembly.*

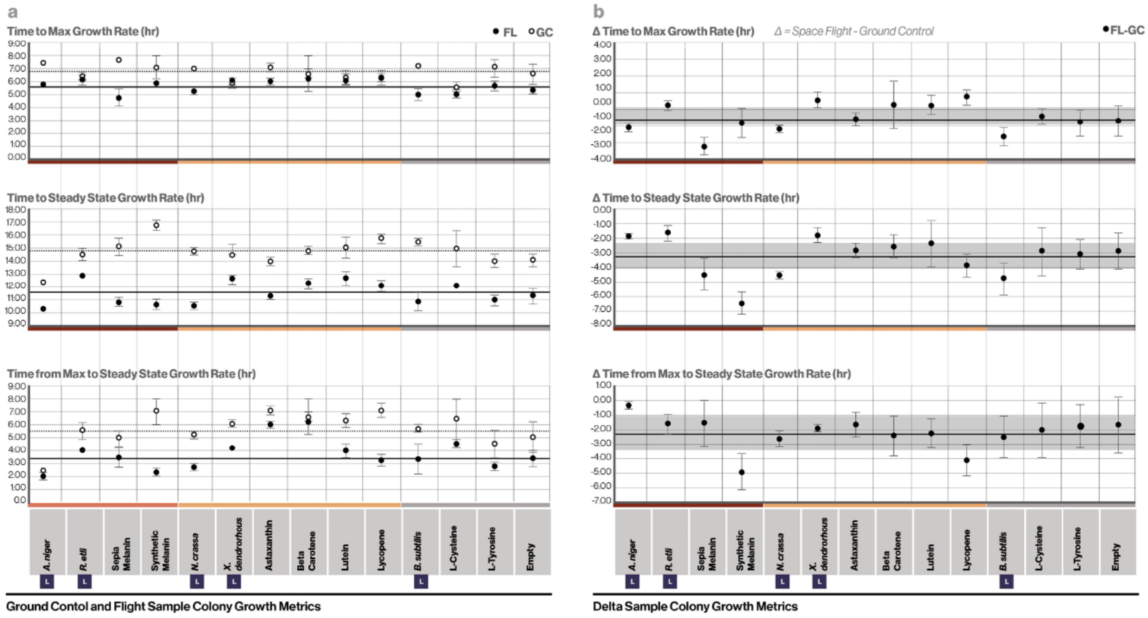


*Supplementary Figure 2. Detailed growth analysis studies. (a) Colony growth metrics for each sample type. Error bars represent standard error of the mean. Horizontal lines represent the mean values across pigment groupings Melain, Carotenoid, and No Pigment, from left to right, for FL (solid line) and GC (dashed line) samples. (b) Deltas between FL and GC samples for each sample type. Error bars represent standard error of the mean. Horizontal lines represent the mean delta value across all samples(solid line), and the shaded region superimposed on the line represents the standard mean error of the delta value across all samples (gray).*
